# Contextual influence on confidence judgments in human reinforcement learning

**DOI:** 10.1101/339382

**Authors:** Maël Lebreton, Karin Bacily, Stefano Palminteri, Jan B. Engelmann

## Abstract

The ability to correctly estimate the probability of one’s choices being correct is fundamental to optimally re-evaluate previous choices or to arbitrate between different decision strategies. Experimental evidence nonetheless suggests that this metacognitive process -referred to as a confidence judgment-is susceptible to numerous biases. We investigate the effect of outcome valence (gains or losses) on confidence while participants learned stimulus-outcome associations by trial-and-error. In two experiments, we demonstrate that participants are more confident in their choices when learning to seek gains compared to avoiding losses. Importantly, these differences in confidence were observed despite objectively equal choice difficulty and similar observed performance between those two contexts. Using computational modelling, we show that this bias is driven by the context-value, a dynamically updated estimate of the average expected-value of choice options that has previously been demonstrated to be necessary to explain equal performance in the gain and loss domain. The biasing effect of context-value on confidence, also recently observed in the context of incentivized perceptual decision-making, is therefore domain-general, with likely important functional consequences.

## Introduction

Simple reinforcement learning algorithms efficiently learn by trial-and-error to implement decision policies that maximize the occurrence of rewards and minimize the occurrence of punishments (Sutton and Barto, 1998). Such basic algorithms have been extensively used in experimental psychology, neuroscience and economics, and seem to parsimoniously account for a large amount of experimental data at the behavioral (Erev and Roth, 1998; Rescorla and Wagner, 1972) and neuronal levels (Daw et al., 2006; O’Doherty et al., 2004; Schultz et al., 1997), as well as for learning abnormalities due to specific pharmacological manipulations (Frank et al., 2004; Pessiglione et al., 2006) and neuro-psychiatric disorders (Palminteri et al., 2012). Yet, ecological environments are inherently ever-changing, volatile and complex, such that organisms need to be able to flexibly adjust their learning strategies or to dynamically select among different learning strategies. These more sophisticated behaviors can be implemented by reinforcement-learning algorithms which compute different measures of environmental uncertainty (Courville et al., 2006; Mathys et al., 2011; Yu and Dayan, 2005) or strategy reliability (Collins and Koechlin, 2012; Daw et al., 2005; Doya et al., 2002). To date, surprisingly little research has investigated if and how individuals engaged in learning by trial-and-error can actually compute such reliability estimates or related proxy variables. One way to experimentally assess such reliability estimates is via eliciting confidence judgments. Confidence is defined as a decision-maker’s estimation of her probability of being correct (Adams, 1957; Pouget et al., 2016; Sanders et al., 2016). It results from a meta-cognitive operation (Fleming and Dolan, 2012), which according to recent studies could be performed automatically even when confidence judgments are not explicitly required (Lebreton et al., 2015). In the context of predictive-inference tasks, individuals’ subjective confidence judgments have been shown to track the likelihood of decisions being correct in changing environments with remarkable accuracy (Heilbron and Meyniel, 2018; Meyniel et al., 2015a). Confidence could therefore be employed as a meta-cognitive variable that enables dynamic comparisons of different learning strategies and ultimately, decisions about whether to adjust learning strategies. Despite the recent surge of neural, computational and behavioral models of confidence estimation in decision-making and prediction tasks (Fleming and Daw, 2017; Meyniel et al., 2015b; Pouget et al., 2016), how decision-makers estimate their confidence in their choices in reinforcement-learning contexts remains poorly investigated.

Crucially, although confidence judgments have been reported to accurately track decision-makers probability of being correct (Meyniel et al., 2015a; Sanders et al., 2016), they are also known to be subject to various biases. Notably, it appears that individuals are generally overconfident regarding their own performance (Lichtenstein et al., 1982), and that confidence judgments are modulated by numerous psychological factors including desirability biases (Giardini et al., 2008), arousal (Allen et al., 2016), mood (Koellinger and Treffers, 2015), and emotions (Jönsson et al., 2005) such as anxiety (Massoni, 2014). Given the potential importance of confidence in mediating learning strategies in changing environments, investigating confidence and confidence biases in reinforcement-learning appears crucial.

Here, following the recent demonstration that confidence in a decision is biased by the value at stake in a perceptual decision-making task (Lebreton et al., 2018), we simultaneously investigated the learning behavior and confidence estimations of individuals engaged in a reinforcement-learning task where the valence of the decision outcomes was systematically manipulated (gains versus losses) (Palminteri et al., 2015; Pessiglione et al., 2006). We hypothesized that individuals would exhibit lower confidence in their choices while learning to avoid losses compared to seeking gains, despite similar performance and objectively equal difficulty between these two learning contexts. In addition, we anticipated that this bias would be generated by the learned context-value associated with decision states (Klein et al., 2017; Palminteri et al., 2015).

Our results, which confirm this hypothesis, illustrate the generalizability of the confidence bias induced by the valence of incentives and outcomes (Lebreton et al., 2018), and suggest that –despite apparent similar behavior-profound asymmetries might exists between learning to avoid losses and learning to seek gains (Palminteri and Pessiglione, 2017), with likely important functional consequences.

## Results

### Experiment 1

We invited 18 participants to partake in our first experiment, and asked them to perform a probabilistic instrumental-learning task adapted from a previous study (Palminteri et al., 2015, 2016). Participants repeatedly faced pairs of abstract symbols probabilistically associated with monetary outcomes. Symbol pairs were fixed, and associated with two levels of two outcome features, namely valence and information, in a 2×2 factorial design. Therefore, pairs of symbols could be associated with either gains or losses, and with partial or complete feedback (**Methods** and Figure 1.A-B). Participants could maximize their payoffs by learning to choose the most advantageous symbol of each pair, i.e., the highest expected gain or the lowest expected loss. At each trial, after their choice but before receiving feedback, participants were also asked to report their confidence in their choice on a Likert scale from 0 to 10. Replicating previous findings (Palminteri et al., 2015, 2016), we found that participants correctly learned by trial-and-error to choose the best outcomes, (average correct choice rate 76.50 ± 2.38, t-test vs chance t_17_ = 11.16; P = 3.04×10^−9^), and that learning performance was marginally affected by the information factor, but unaffected by the outcome valence (ANOVA; main effect of information F_1,17_ = 4.28; P = 0.05; main effect of valence F_1,17_ = 1.04; P = 0.32; interaction F_1,17_ = 1.06; P = 0.32; Figure 1.C). In other words, participants learned equally well to seek gains and to avoid losses. However, and in line with our hypothesis, the confidence ratings showed a very dissimilar pattern, as they were strongly influenced by the valence of outcomes (ANOVA; main effect of information F_1,17_ = 2.00; P = 0.17; main effect of valence F_1,17_ = 33.11; P = 2,33×10^−11^; interaction F_1,17_ = 7.58; P = 0.01; Figure 1.D). Similar to the valence bias reported in perceptual decision-making tasks (Lebreton et al., 2018), these effects were driven by the fact that participants were more confident in the gain than in the loss condition when receiving partial feedback (6.86 ± 0.28 vs 4.66 ± 0.39; t-test t_17_ = 7.20; P = 1.50×10^−6^), and that this difference was still very significant although smaller in the complete feedback condition (6.58 ± 0.35 vs 5.24 ± 0.37; t-test t_17_ = 3.52; P = 2.65×10^−3^).

**Figure 1.**
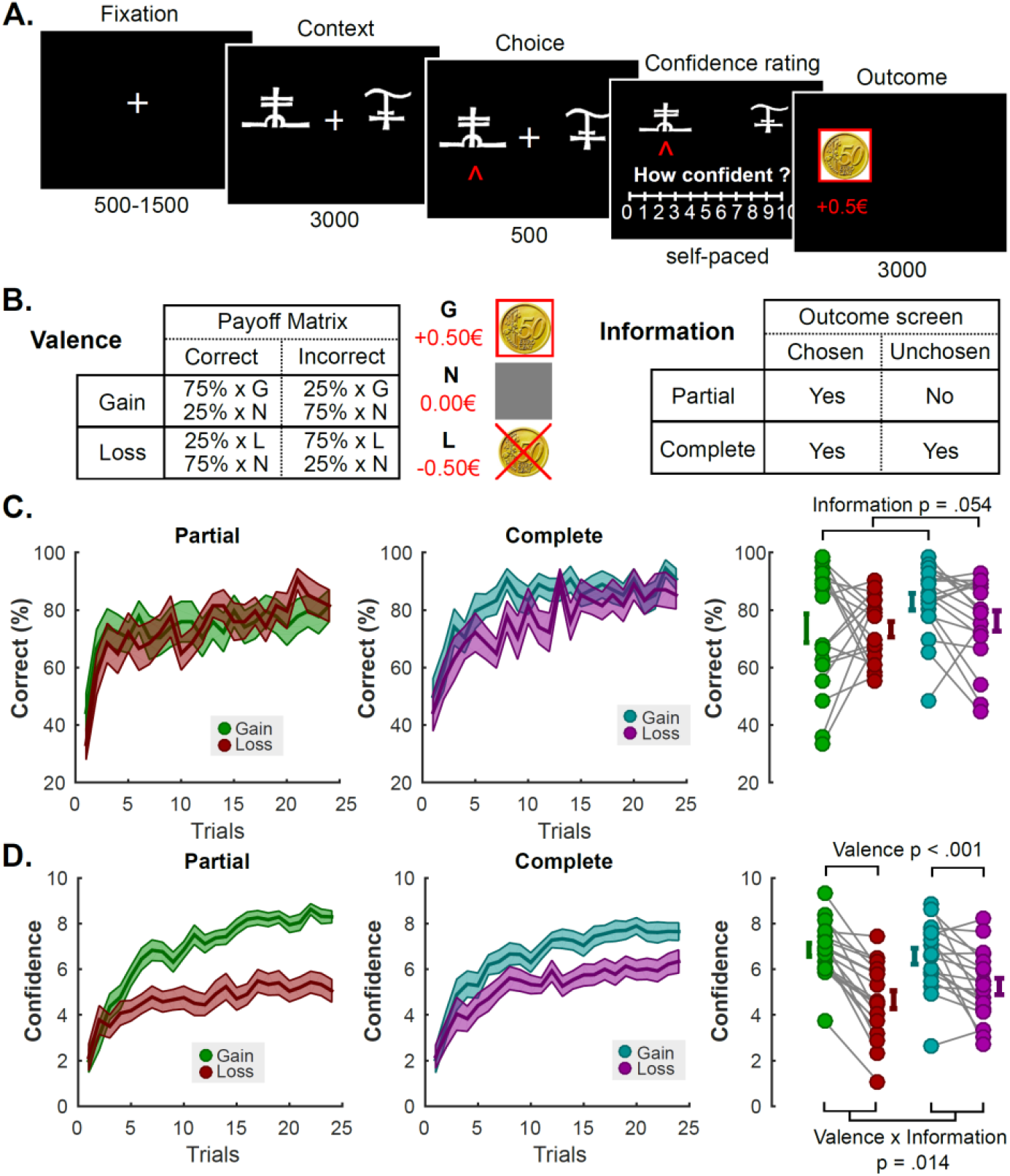
Experiment 1 Task Schematic, Learning and Confidence Results. A. **Behavioral task**. Successive screens displayed in one trial are shown from left to right with durations in ms. After a fixation cross, participants viewed a couple of abstract symbols displayed on both sides of a computer screen and had to choose between them. They were thereafter asked to report their confidence in their choice on a numerical scale (graded from 0 to 10). Finally, the outcome associated with the chosen symbol was revealed.
B. **Task design and contingencies**.
C. **Performance**. Trial by trial percentage of correct responses in the partial (left) and the complete (middle) information conditions. Filled colored areas represent mean ± sem; Right: Individual averaged performances in the different conditions. Connected dots represent individual data points in the within-subject design. The error bar displayed on the side of the scatter plots indicate the sample mean ± sem.
D. **Confidence**. Trial by trial confidence ratings in the partial (left) and the complete (middle) information conditions. Filled colored areas represent mean ± sem; Right: Individual averaged performances in the different conditions. Connected dots represent individual data points in the within-subject design. The error bar displayed on the side of the scatter plots indicate the sample mean ± sem.

### Experiment 2

While the results of the first experiment are strongly suggestive of an effect of outcome valence on confidence in reinforcement learning, they cannot *formally* characterize a bias, as the notion of cognitive bias depends on the optimal reward-maximizing strategy (Marshall et al., 2013). In other terms: does this bias persist in situations where a truthful and accurate confidence report is associated with payoff maximization? We addressed this limitation of experiment 1 by directly incentivizing reports of confidence accuracy in our follow-up experiment. In this new experiment, confidence was formally defined as an estimation of the probability of being correct, and participants could maximize their chance to gain an additional monetary bonus (3×5 euros) by reporting their confidence as accurately and truthfully as possible on a rating scale ranging from 50% to 100% (Figure 2.A). Specifically, confidence judgments were incentivized with a Matching Probability (MP) mechanism, a well-validated method from behavioral economics adapted from the Becker-DeGroot-Marschak auction (Becker et al., 1964; Ducharme and Donnell, 1973). Briefly, the MP mechanism considers participants’ confidence reports as bets on the correctness of their answers, and implements comparisons between these bets and random lotteries (Figure 3.A). Under utility maximization assumptions, this guarantees that participants maximize their earnings by reporting their most precise and truthful confidence estimation (Schlag et al., 2015; Schotter and Trevino, 2014). This mechanism and the dominant strategy were explained to the 18 new participants before the experiment (**Methods**). In addition, because the neutral and non-informative outcome was more frequently experienced in the punishment partial than in the reward partial context in experiment 1, we replaced the neutral 0€ with a 10c gain or loss (see **Methods** and Figure 2.B).

**Figure 2.**
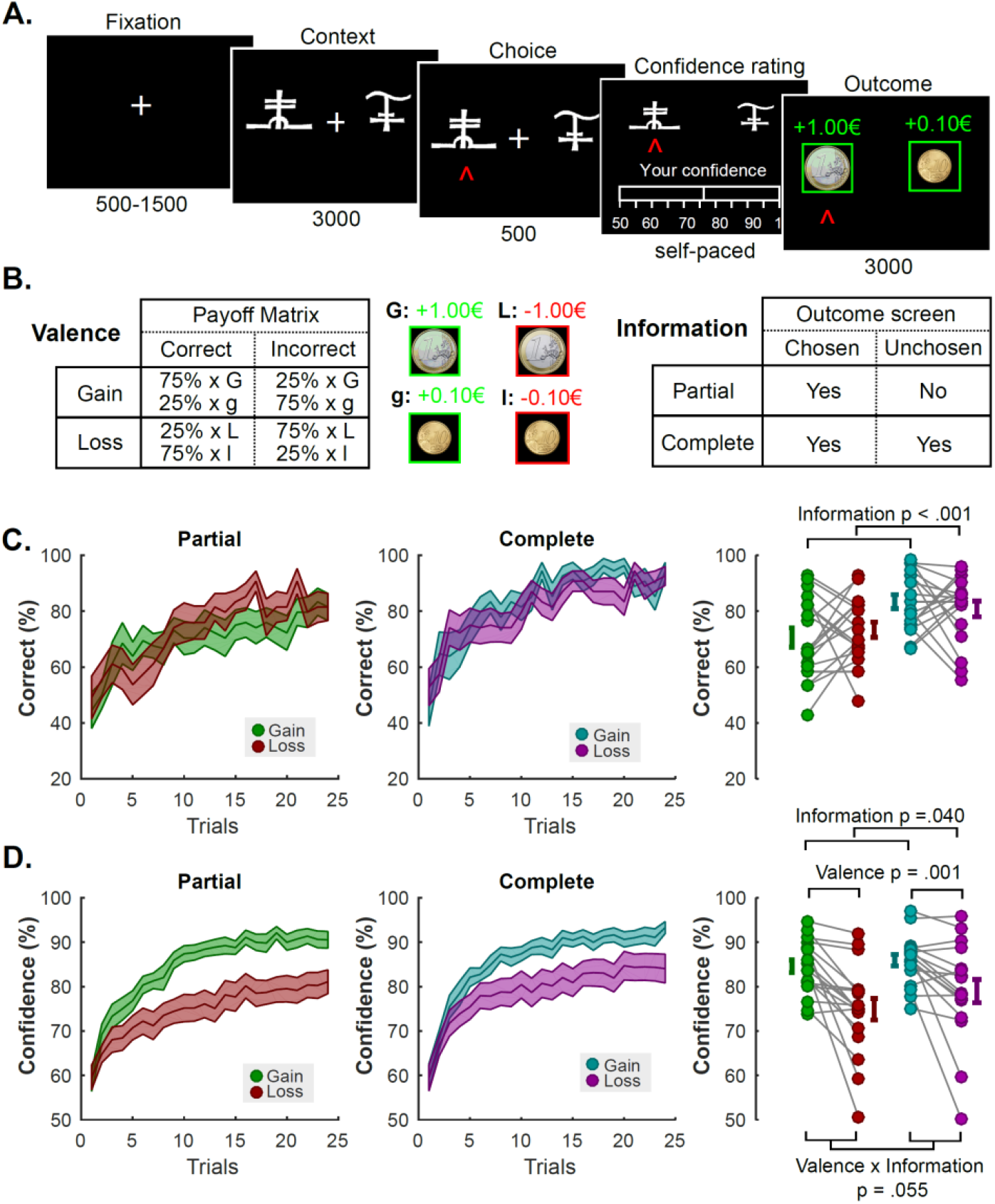
Experiment 2 Task Schematic, Learning and Confidence Results. A. **Behavioral task**. Successive screens displayed in one trial are shown from left to right with durations in ms. After a fixation cross, participants viewed a couple of abstract symbols displayed on both sides of a computer screen, and had to choose between them. They were thereafter asked to report their confidence in their choice on a numerical scale (graded from 50 to 100%). Finally, the outcome associated with the chosen symbol was revealed.
B. **Task design and contingencies**.
C. **Performance**. Trial by trial percentage of correct responses in the partial (left) and the complete (middle) information conditions. Filled colored areas represent mean ± sem; Right: Individual averaged performances in the different conditions. Connected dots represent individual data points in the within-subject design. The error bar displayed on the side of the scatter plots indicate the sample mean ± sem.
D. **Confidence**. Trial by trial confidence ratings in the partial (left) and the complete (middle) information conditions. Filled colored areas represent mean ± sem; Right: Individual averaged performances in the different conditions. Connected dots represent individual data points in the within-subject design. The error bar displayed on the side of the scatter plots indicate the sample mean ± sem.

**Figure 3.**
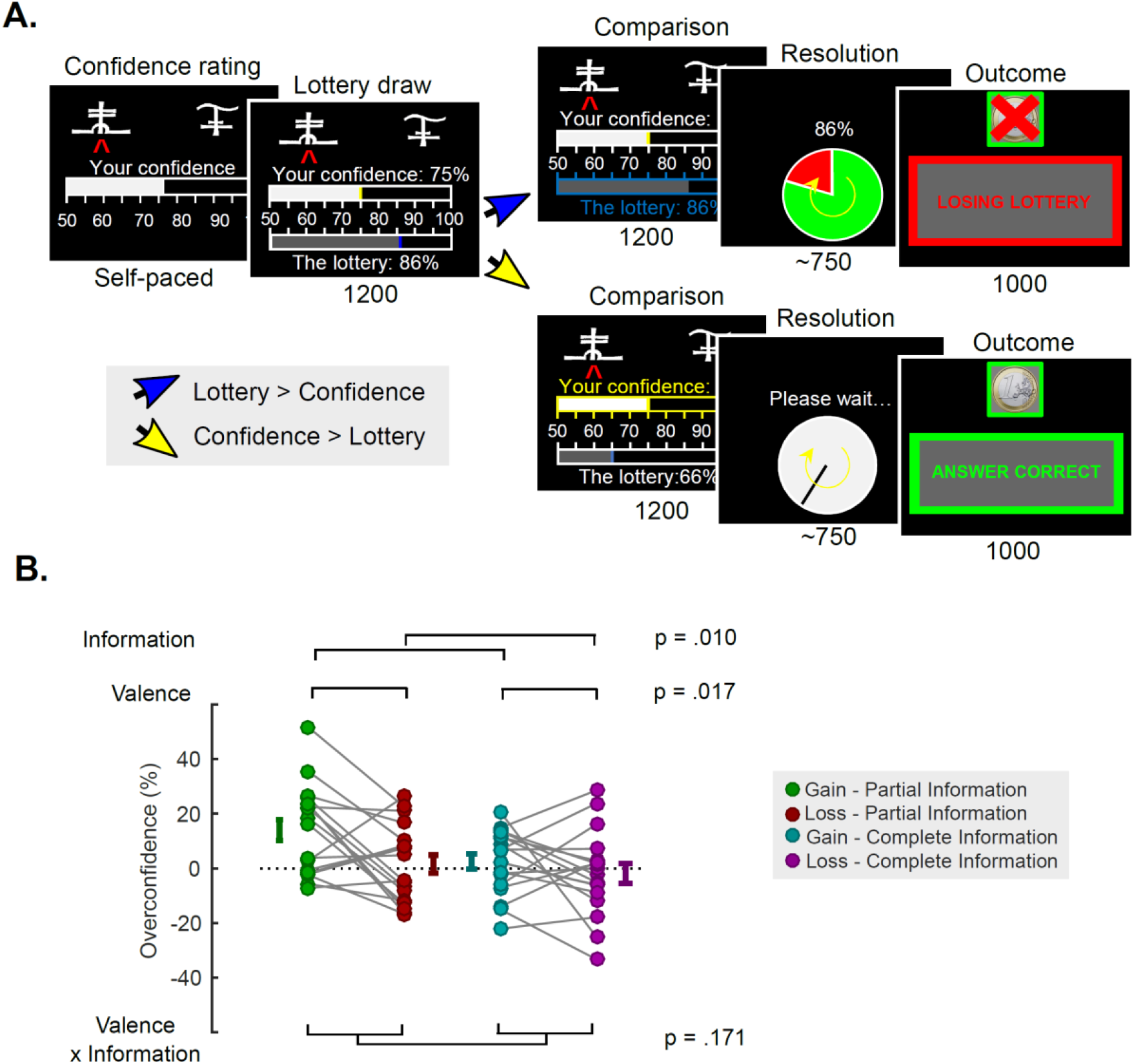
Incentive mechanism and overconfidence. A. **Incentive mechanism**. In Experiment 2, for the payout-relevant trials a lottery L is randomly drawn in the 50-100% interval and compared to the confidence rating C. If L > C, the lottery is implemented. A wheel of fortune, with a L% chance of losing is displayed, and played out. Then, feedback informed participants whether the lottery resulted in a win or a loss. If C > L, a clock is displayed together with the message “Please wait”, followed by feedback which depended on the correctness of the initial choice. With this mechanism, participant can maximize their earning by reporting their confidence accurately and truthfully.
B. **Overconfidence**. Individual averaged calibration, as a function of Experiment 2 experimental conditions (with a similar color code as in Figure 1–2). Connected dots represent individual data points in the within-subject design. The error bar displayed on the side of the scatter plots indicate the sample mean ± sem.

Replicating the results from the first experiment, we found that learning performance was affected by the information factor, but unaffected by the outcome valence (ANOVA; main effect of information F_1,17_ = 18.64; P = 4.67×10^−4^; main effect of valence F_1,17_ = 1.33×10^−3^; P = 0.97; interaction F_1,17_ = 0.77; P = 0.39; Figure 2.C). Yet, the confidence ratings were again strongly influenced by the valence of outcomes (ANOVA; main effect of information F_1,17_ = 4.92; P = 0.04; main effect of valence F_1,17_ = 15.43; P = 1.08×10^−3^; interaction F_1,17_ = 4.25; P = 0.05; Figure 2.D). Similar to Experiment 1, these effects were driven by the fact that participants were more confident in the gain than in the loss conditions (85.25 ± 1.23 vs 76.96 ± 2.38 (in %); t-test t_17_ = 3.93; P = 1.08×10^−3^).

Importantly, the changes in the experimental design also allowed us to estimate the bias in confidence judgments (sometimes called calibration, or “overconfidence”), by contrasting individuals’ average reported confidence (i.e. estimated probability of being correct) with their actual average probability of being correct. A positive bias therefore indicates that participants are overconfident reporting a higher probability of being correct than their objective average performance. Conversely, a negative bias indicates reporting a lower probability of being correct than the true average (“underconfidence”). These analyses revealed that participants are, in general marginally overconfident (4.07 ± 2.37 (%); t-test vs 0: t_17_ = 1.72; P = 0.10). This overconfidence, which was maximal in the gain-partial information condition (14.00 ± 3.86 (%)), was nonetheless mitigated by complete information (gain-complete: 2.53 ± 2.77 (%); t-test vs gain-partial: t_17_ = 2.72; P = 0.01) and losses (loss-partial: 1.56 ± 3.35 (%); t-test vs gain-partial: t_17_ = 2.76; P = 0.01). These effects of outcome valence and counterfactual feedback information on overconfidence appeared to be simply additive (ANOVA; main effect of information F_1,17_ = 8.40; P = 0.01; main effect of valence F_1,17_ = 7.03; P = 0.02; interaction F_1,17_ = 2.05; P = 0.17; Figure 3.B).

### Context-dependent learning

While the results from our two first experiments provide convincing support for our hypotheses at the aggregate level (i.e. averaged choice rate and confidence ratings), we aimed at providing a finer description of the dynamical processes at stake, and therefore turned to computational modelling. Standard reinforcement-learning algorithms (Rescorla and Wagner, 1972; Sutton and Barto, 1998) typically give a satisfactory account of learning dynamics in stable contingency tasks as ours, but recent studies (Klein et al., 2017; Palminteri et al., 2015, 2016) have demonstrated that human learning is highly context (or reference)-dependent. This context dependency, by allowing neutral or moderately negative outcomes to be reframed as relative gains, provides an effective and parsimonious solution to the punishment-avoidance paradox. In addition, context dependency accounts for “irrational choices” observed in a transfer task performed after learning: participants express higher preference for mildly unfavorable items to objectively better items, because the former were initially paired with unfavorable items and hence acquired a higher “relative” subjective value (Klein et al., 2017; Palminteri et al., 2015, 2016). As in these previous studies, the participants from our two experiments also performed the transfer test after the learning task. The typical behavioral signature of context-dependent learning is a preference reversal in the complete information contexts, where symbols associated with small losses (L_25_) becomes preferred to symbols associated with small gains (G_25_), despite having objectively lower expected value (Klein et al., 2017; Palminteri et al., 2015, 2016). This pattern was present in both our experiments (% choices; experiment 1: L_25_: 59.52 ± 4.88, G_25_: 38.89 ± 5.04; t-test t_17_ = 2.46; P = 0.02; experiment 2: L_25_: 67.26 ± 5.35, G_25_: 28.37 ± 4.46; t-test t_17_ = 5.27; P = 6.24×10^−5^, see Figure 4 A-B, middle panels).

To confirm these observations, we adopted a model-fitting and model-comparison approach, where a standard learning model (ABSOLUTE) was compared to context-dependent learning model (RELATIVE) in their ability to account for the participant choices (**Methods**). Replicating previous findings (Palminteri et al., 2015, 2016), the context-dependent model provided the best and most parsimonious account of the data collected in our 2 experiments (Table 1), and a satisfactory account of choice patterns in both the learning (average likelihood per trial in experiment 1: 0.72 ± 0.03; in experiment 2: 0.72 ± 0.02; see Figure 4 A-B, top panels) and transfer tasks (average likelihood per trial; experiment 1: 0.71 ± 0.02; experiment 1: 0.70 ± 0.02; see Figure 4 A-B, middle panels). Please also note that the model estimated free-parameters (Table 2) are very similar to what was reported in the previous studies (Palminteri et al., 2015, 2016).

**Figure 4.**
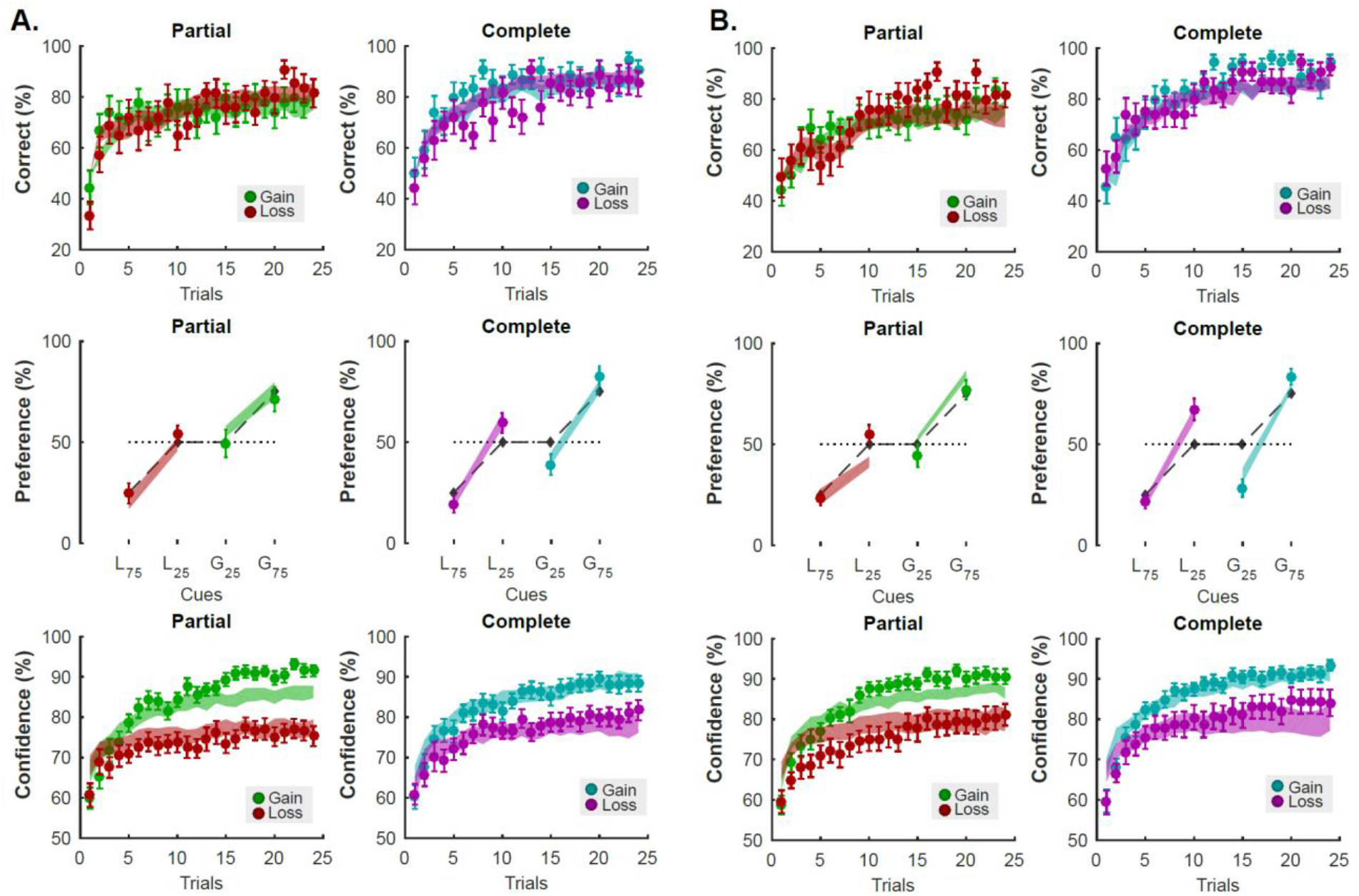
Modelling results: fits. Behavioral results and model fits in Experiments 1(**A**) and 2 (**B**). Top: Learning performance (i.e. percent correct). Middle: Choice rate in the transfer test. Symbols are ranked by expected value (L_75_: symbol associated with 75% probability of losing 1€; L_25_: symbol associated with 25% probability of losing 1€; G_25_: symbol associated with 25% probability of winning 1€; G_75_: symbol associated with 75% probability of winning 1€;) Bottom: Confidence ratings. In all panels, colored dots and error bars represent the actual data (mean ± sem), and filled areas represent the model fits (mean ± sem). Model fits were obtained with the RELATIVE reinforcement learning model for the learning performance (top) and the choice rate in the transfer test (middle), and with the FULL glme for the confidence ratings (bottom). Dark grey diamonds in the Preference panels (middle) indicate the expected preference probability given the symbols objective expected value (L_75_: −0.75€; L_25_: −0.25€; G_25_: 0.25€; G_75_: 0.75€);

**Table 1.** Reinforcement-learning. Model comparison. AIC, Akaike Information Criterion (computed with LLmax); BIC, Bayesian Information Criterion (computed with LLmax); DF, degrees of freedom; LLmax, maximal log likelihood; LPP, log of posterior probability; PP, posterior probability of the model given the data; XP, exceedance probability (computed from LPP). The table summarizes for each model its fitting performances.

|  |  |  |  |  |  |  |  |  |
| --- | --- | --- | --- | --- | --- | --- | --- | --- |
| Exp. 1 | **Model** | **DF** | **-2\*LLmax** | **2\*AIC** | **BIC** | **-2\*LPP** | **EF** | **XP** |
|  | ABSOLUTE | 3 | 385±20 | 392±20 | 404±20 | 391±20 | 0.12 | 0.0 |
|  | RELATIVE | 4 | 345±24 | 353±24 | 369±24 | 354±24 | 0.88 | 1.0 |
| Exp. 2 | **Model** | **DF** | **-2\*LLmax** | **2\*AIC** | **BIC** | **-2\*LPP** | **EF** | **XP** |
|  | ABSOLUTE | 3 | 411±15 | 417±15 | 429±15 | 416±15 | 0.05 | 0.0 |
|  | RELATIVE | 4 | 355±16 | 363±16 | 379±16 | 362±16 | 0.95 | 1.0 |

**Table 2.** Reinforcement-learning. Free parameters. ABSOLUTE, absolute value learning model; RELATIVE, relative value learning model (best-fitting model); LL maximization, parameters obtained when maximizing the negative log likelihood; LPP maximization, parameters obtained when maximizing the negative log of the Laplace approximation of the posterior probability. The table summarizes for each model the likelihood maximizing (best) parameters averaged across subjects. Data are expressed as mean±s.e.m. The values retrieved from the LPP maximization procedure are those used to generate the variable used in the confidence glme models.

|  |  | LL Maximization |  | LPP Maximization |  |
| --- | --- | --- | --- | --- | --- |
| Exp. 1 | Free Parameter | ABSOLUTE | RELATIVE | ABSOLUTE | RELATIVE |
| | Inverse temperature ($\beta$) | 6.29±0.63 | 54.04±38.8 | 6.07±0.61 | 12.65±1.47 |
| | Factual learning rate ($\alpha_c$) | 0.37±0.05 | 0.23±0.04 | 0.36±0.04 | 0.24±0.04 |
| | Counterfactual learning rate ($\alpha_u$) | 0.13±0.03 | 0.07±0.02 | 0.15±0.03 | 0.09±0.02 |
| | Context learning rate ($\alpha_v$) | - | 0.46±0.10 | - | 0.46±0.10 |
|  |  | LL Maximization |  | LPP Maximization |  |
| Exp. 2 | Free Parameter | ABSOLUTE | RELATIVE | ABSOLUTE | RELATIVE |
| | Inverse temperature ($\beta$) | 102.00±99.49 | 83.05±73.15 | 2.65±0.29 | 6.86±0.81 |
| | Factual learning rate ($\alpha_c$) | 0.49±0.07 | 0.26±0.04 | 0.49±0.07 | 0.24±0.04 |
| | Counterfactual learning rate ($\alpha_u$) | 0.24±0.08 | 0.12±0.04 | 0.24±0.08 | 0.13±0.03 |
| | Context learning rate ($\alpha_v$) | - | 0.41±0.09 | - | 0.40±0.09 |

### The model-free and model-based determinants of confidence and performance

We next used latent variables from this computational model, along with other variables known to inform confidence judgments, to inform a descriptive model of confidence formation. We propose confidence to be under the influence of three main variables, entered as explanatory variables in linear mixed-effect regressions (FULL model). The first explanatory variable is choice difficulty, a feature captured in value-based choices by the absolute difference between the expected value of the two choice options (De Martino et al., 2013; Folke et al., 2016), and indexed by the absolute difference between the option Q-values calculated by the RELATIVE model. The second explanatory variable is the confidence expressed at the preceding trial. Confidence judgments indeed exhibit a strong auto-correlation, even when they relate to decisions made in different tasks (Rahnev et al., 2015). Note that in our task, where the stimuli are presented in an interleaved design, this last term captures the features of confidence which are transversal to different contexts such as aspecific drifts due to attention fluctuation and/or fatigue. The third and final explanatory variable is V(s), the approximation of the average expected-value of a pair of stimuli (i.e., the context value from the RELATIVE model) (Palminteri et al., 2015). The context value, initialized at zero, gradually becomes positive in the reward-seeking conditions and negative in the punishment-avoidance conditions. This variable is central to our hypothesis that the decision frame (gain vs. loss) influences individuals’ estimated confidence about being correct (Lebreton et al., 2018). Crucially, in the FULL model, all included explanatory variables were significant predictors of confidence ratings in both experiments (see Table 3). As a quality check, we also verified that the confidence ratings estimated under the FULL model satisfactorily capture the evolution of observed confidence ratings across the course of our experiments (Figure 4 A-B, bottom panels).

**Table 3.** Modelling confidence ratings. Estimated fixed-effect coefficients from generalized linear mixed-effect models.

|  |  | GLME |  |
| --- | --- | --- | --- |
| Experiment 1 | Fixed-Effect | REDUCED | FULL |
| | Intercept ($\beta_0$) | 0.52±0.04<br>$t_{5079} = 14.46$; $P = 1.90 \times 10^{-46}$ | 0.53±0.04<br>$t_{5078} = 14.55$; $P = 4.92 \times 10^{-47}$ |
| | Choice difficulty ($\beta_{\Delta Q}$) | 0.33±0.06<br>$t_{5079} = 5.77$; $P = 8.43 \times 10^{-9}$ | 0.30±0.05<br>$t_{5078} = 5.96$; $P = 2.73 \times 10^{-9}$ |
| | Preceding confidence ($\beta_{c1}$) | 0.28±0.04<br>$t_{5079} = 7.60$; $P = 3.62 \times 10^{-14}$ | 0.28±0.03<br>$t_{5078} = 7.39$; $P = 1.67 \times 10^{-13}$ |
| | Context value ($\beta_V$) | - | 0.47±0.14<br>$t_{5078} = 3.21$; $P = 1.35 \times 10^{-3}$ |
|  |  | GLME |  |
| Experiment 2 | Fixed-Effect | REDUCED | FULL |
| | Intercept ($\beta_0$) | 0.53±0.03<br>$t_{5145} = 17.57$; $P = 3.77 \times 10^{-67}$ | 0.53±0.03<br>$t_{5144} = 17.12$; $P = 5.94 \times 10^{-64}$ |
| | Choice difficulty ($\beta_{\Delta Q}$) | 0.18±0.02<br>$t_{5145} = 6.33$; $P = 2.63 \times 10^{-10}$ | 0.17±0.03<br>$t_{5144} = 5.90$; $P = 3.85 \times 10^{-9}$ |
| | Preceding confidence ($\beta_{c1}$) | 0.29±0.04<br>$t_{5145} = 7.01$; $P = 2.75 \times 10^{-12}$ | 0.30±0.04<br>$t_{5144} = 7.48$; $P = 8.54 \times 10^{-14}$ |
| | Context value ($\beta_V$) | - | 0.16±0.06<br>$t_{5144} = 2.51$; $P = 1.19 \times 10^{-2}$ |

On the contrary, when attempting to predict the trial-by-trial correct answers (i.e. performance) rather than confidence judgments with the same explanatory variables, the choice difficulty and the confidence expressed at the preceding trial were significant predictors in the two experiments, while the context value was not (Table 4). This again captures the idea that the context value might bias confidence judgments, above and beyond the variation in performance. Finally, because decision reaction times are known to be (negatively) correlated with subsequent confidence judgments - the more confident individuals are in their choices, the faster their decisions (De Martino et al., 2013; Kiani et al., 2014; Lebreton et al., 2015)-, we anticipated and verified that the same explanatory variables which are significant predictors of confidence also predict reaction times (although with opposite signs – see Table 4).

**Table 4.** Modelling performance and reaction times. Estimated fixed-effect coefficients from generalized linear mixed-effect models (performance: logistic regression; reaction times: linear regression).

|  |  | GLME |  |
| --- | --- | --- | --- |
| Experiment 1 | Fixed-Effect | PERFORMANCE | RT |
| | Intercept ($\beta_0$) | -0.84±0.20<br>$t_{5078} = -4.15$; $P = 3.40 \times 10^{-5}$ | 1.90±0.09<br>$t_{5078} = 20.12$; $P = 1.12 \times 10^{-86}$ |
| | Choice difficulty ($\beta_{\Delta Q}$) | 9.90±1.67<br>$t_{5078} = 5.92$; $P = 3.32 \times 10^{-9}$ | -0.65±0.20<br>$t_{5078} = -3.15$; $P = 1.63 \times 10^{-3}$ |
| | Preceding confidence ($\beta_{c1}$) | 1.28±0.36<br>$t_{5078} = 3.60$; $P = 3.19 \times 10^{-4}$ | -0.24±0.14<br>$t_{5078} = -1.78$; $P = 0.08$ |
| | Context value ($\beta_V$) | 1.19±0.54<br>$t_{5078} = 2.19$; $P = 0.03$ | -0.37±0.11<br>$t_{5078} = -3.48$; $P = 5.04 \times 10^{-4}$ |
|  |  | GLME |  |
| Experiment 2 | Fixed-Effect | PERFORMANCE | RT |
| | Intercept ($\beta_0$) | -0.71±0.22<br>$t_{5144} = -3.20$; $P = 1.37 \times 10^{-3}$ | 1.68±0.09<br>$t_{5144} = 17.93$; $P = 9.09 \times 10^{-70}$ |
| | Choice difficulty ($\beta_{\Delta Q}$) | 5.29±0.76<br>$t_{5144} = 6.94$; $P = 4.49 \times 10^{-12}$ | -0.41±0.09<br>$t_{5144} = -4.50$; $P = 6.81 \times 10^{-6}$ |
| | Preceding confidence ($\beta_{c1}$) | 1.21±0.33<br>$t_{5144} = 3.66$; $P = 2.57 \times 10^{-4}$ | -0.54±0.10<br>$t_{5144} = -5.31$; $P = 1.08 \times 10^{-7}$ |
| | Context value ($\beta_V$) | 0.30±0.28<br>$t_{5144} = 1.05$; $P = 0.29$ | -0.17±0.05<br>$t_{5144} = -3.68$; $P = 2.35 \times 10^{-4}$ |

### Context values explain the confidence bias

In this last section, we aimed at demonstrating that the context values are necessary and sufficient to explain the difference in confidence observed between the reward seeking and the loss avoidance conditions. We therefore built a REDUCED model, which was similar to the FULL model, but lacked the context value (see Table 3). First, because the REDUCED model is nested in the FULL model, a likelihood ratio test statistically assesses the probability of observing the estimated fitting difference under the null hypothesis that the FULL model is not better than the REDUCED model. In both experiments, this null hypothesis was rejected (both P<0.001), indicating that the FULL model provides a better explanation of the observed data. Hence confidence is critically modulated by the context value.

Then, to demonstrate that the biasing effect of outcome valence on confidence is operated through the context value, we show that the REDUCED model, which lacks the context value as an explanatory variable, cannot reproduce the critical pattern of valence-induced confidence biases observed in our data, while the FULL model can (Figure 5) (Palminteri et al., 2017). Overall, these results provide additional evidence for the importance of context value as an important latent variable in learning, not only explaining irrational choices in transfer tests, but also confidence biases observed during learning (Figure 6).

**Figure 5.**
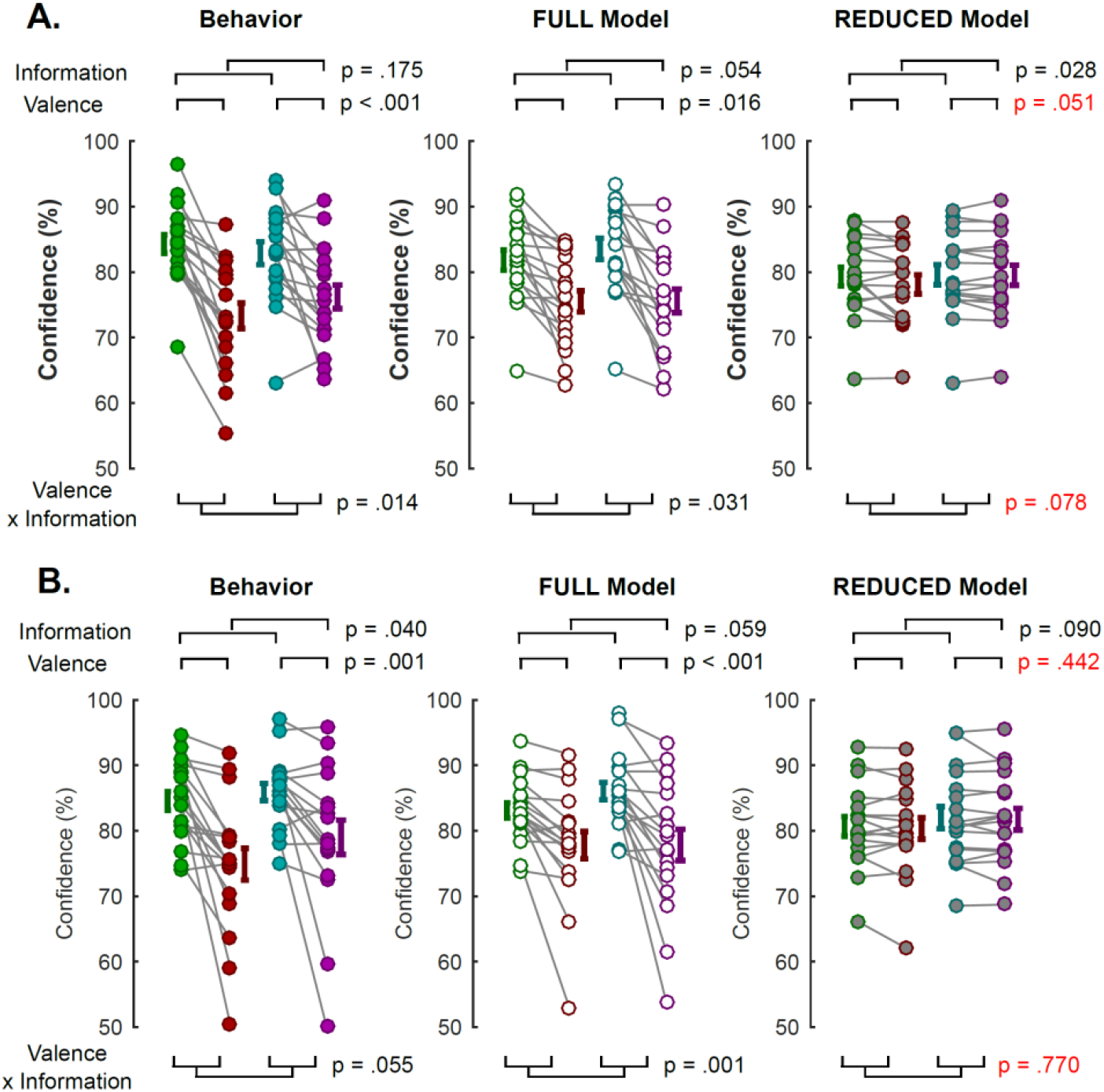
Modelling results: lesioning approach. Two nested models are compared in their ability to reproduce the pattern of interest observed in averaged confidence ratings, in experiment 1 (**A**) and experiment 2 (**B**). In the FULL model, confidence is modelled as a function of three factors: the absolute difference between options values, the confidence observed in the previous trial, and the context value. In the REDUCED model, confidence is modelled as a function of only two factors: the absolute difference between options values and the confidence observed in the previous trial. Hence, the REDUCED model omits the context-value as a predictor of confidence. Left: pattern of confidence ratings observed in the behavioral data. Middle: pattern of confidence ratings estimated from the FULL model. Right: pattern of confidence ratings estimated from the REDUCED model. In red are reported statistics from a repeated-measure ANOVA where the alternative model fails to reproduce important statistical properties of confidence observed in the data. Connected dots represent individual data points in the within-subject design. The error bar displayed on the side of the scatter plots indicate the sample mean ± sem.

**Figure 6.**
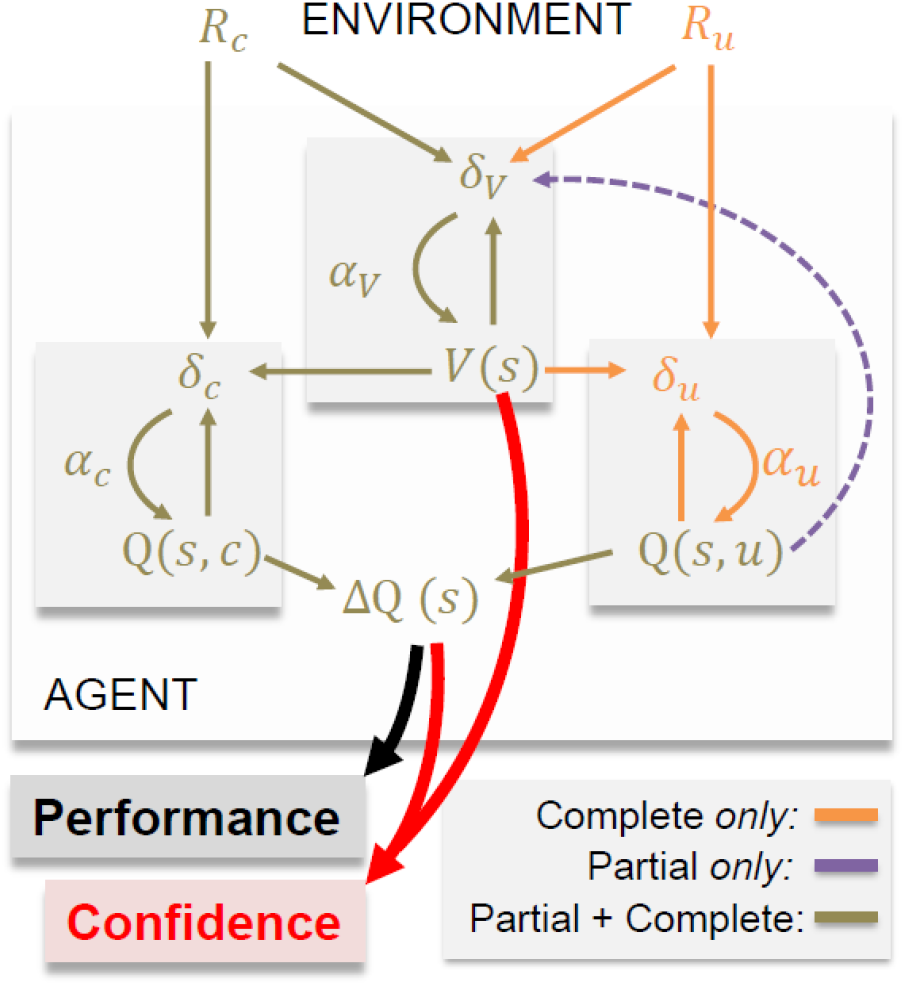
Summary of the modelling results. The schematic illustrates the computational architecture that best account for the choice and confidence data. In each context (or state) ‘s’, the agent tracks option values (Q(s,:)), which are used to decide amongst alternative courses of action, together with the value of the context (V(s)), which quantify the average expected value of the decision context. In all contexts, the agent receives an outcome associated with the chosen option (R_c_), which is used to update the chosen option value (Q(s,c)) via a prediction error (δ_c_) weighted by a learning rate (α_c_). In the complete feedback condition, the agent also receive information about the outcome of the unselected option (R_u_), which is used to update the unselected option value (Q(s,u)) via a prediction error (δ_u_) weighted by a learning rate (α_u_). The available feedback information (R_c_ and R_u_, in the complete feedback contexts and Q(s,u) in the partial feedback contexts) is also used to update the value of the context (V(s)), via a prediction error (δ_v_) weighted by a specific learning rate (α_v_).

### Assessing the specific role of context values in biasing confidence

So far, our investigations show that including context values (V(s)) as a predictor of confidence is necessary and sufficient to reproduce the bias in confidence induced by the decision frame (gain vs. loss). However, it remains unclear how specific and robust the contribution of context-values in generating this bias is, notably when other valence-sensitive model-free and model-based variable are accounted for. To address this question, we run two additional linear models: one including the sum of the two q-values (∑Q), which also track aspects of the valence of the context; the second including RTs, which were also predicted by both ΔQ and V(s) (see previous paragraph). In both experiments and for both linear models, the residual effect of V(s) on trial-by-trial confidence judgments remained positive and (marginally) significant (see Table 5), thus indicating a specific role of our model-driven estimate of V(s) above and beyond other related variables.

**Table 5.** Assessing the specific role of context values on confidence. Estimated fixed-effect coefficients from generalized linear mixed-effect models.

|  |  | GLME |  |
| --- | --- | --- | --- |
| GLME 1 | Fixed-Effect | Experiment 1 | Experiment 2 |
| | Intercept ($\beta_0$) | 0.58±0.05<br>$t_{5077} = 18.06$; $P = 1.01 \times 10^{-70}$ | 0.68±0.03<br>$t_{5143} = 21.74$; $P = 2.48 \times 10^{-100}$ |
| | Choice difficulty ($\beta_{\Delta Q}$) | 0.27±0.05<br>$t_{5077} = 5.55$; $P = 2.97 \times 10^{-8}$ | 0.13±0.03<br>$t_{5143} = 4.97$; $P = 6.76 \times 10^{-7}$ |
| | Preceding confidence ($\beta_{c1}$) | 0.26±0.03<br>$t_{5077} = 7.56$; $P = 4.79 \times 10^{-14}$ | 0.24±0.04<br>$t_{5143} = 6.93$; $P = 4.69 \times 10^{-12}$ |
| | Context value ($\beta_V$) | 0.43±0.14<br>$t_{5077} = 3.14$; $P = 1.68 \times 10^{-3}$ | 0.15±0.06<br>$t_{5143} = 2.36$; $P = 1.81 \times 10^{-2}$ |
| | Reaction times ($\beta_{RT}$) | -0.03±0.01<br>$t_{5077} = -2.53$; $P = 1.15 \times 10^{-2}$ | -0.09±0.01<br>$t_{5143} = -9.95$; $P = 4.04 \times 10^{-24}$ |
|  |  | GLME |  |
| GLME 2 | Fixed-Effect | Experiment 1 | Experiment 2 |
| | Intercept ($\beta_0$) | 0.53±0.04<br>$t_{5077} = 14.99$; $P = 9.36 \times 10^{-50}$ | 0.53±0.03<br>$t_{5143} = 16.83$; $P = 6.45 \times 10^{-62}$ |
| | Choice difficulty ($\beta_{\Delta Q}$) | 0.24±0.05<br>$t_{5077} = 4.59$; $P = 4.53 \times 10^{-6}$ | 0.14±0.03<br>$t_{5143} = 4.79$; $P = 1.75 \times 10^{-6}$ |
| | Preceding confidence ($\beta_{c1}$) | 0.28±0.04<br>$t_{5077} = 7.50$; $P = 7.30 \times 10^{-14}$ | 0.30±0.04<br>$t_{5143} = 7.70$; $P = 1.60 \times 10^{-14}$ |
| | Context value ($\beta_V$) | 0.10±0.05<br>$t_{5077} = 1.94$; $P = 5.22 \times 10^{-2}$ | 0.06±0.02<br>$t_{5143} = 3.96$; $P = 7.50 \times 10^{-5}$ |
| | q-values sum ($\beta_{\Sigma Q}$) | 0.22±0.09<br>$t_{5077} = 2.43$; $P = 1.52 \times 10^{-2}$ | 0.06±0.02<br>$t_{5143} = 2.65$; $P = 7.98 \times 10^{-3}$ |

## Discussion

In this paper we investigated the effect of context-value on confidence during reinforcement-learning, by combining well-validated tasks: a probabilistic instrumental task with monetary gains and losses as outcomes (Palminteri and Pessiglione, 2017; Palminteri et al., 2015; Pessiglione et al., 2006), and two variants of a confidence elicitation task (Hollard et al., 2015; Schlag et al., 2015): a free elicitation of confidence (experiment 1), and an incentivized elicitation of confidence called matching probability (experiment 2). Behavioral results from two experiments consistently show a clear dissociation of the effect of decision frame on learning performance and confidence judgments: while the valence of decision outcomes (gains vs. losses) had no effect on the learning performance, it significantly impacted subjects’ confidence in the very same choices. Specifically, learning to avoid losses generated lower confidence reports than learning to seek gains regardless of the confidence elicitation methods employed. These results extend prior findings (Lebreton et al., 2018) by demonstrating a biasing effect of incentive valence in a reinforcement learning context. They are also consistent with other decision-making studies reporting that positive psychological factors and states, such as joy or desirability, bias confidence upwards, while negative ones, such as worry, bias confidence downwards (Giardini et al., 2008; Jönsson et al., 2005; Koellinger and Treffers, 2015; Massoni, 2014).

Based on the current design and results, we can rule out two potential explanation for the presence of this confidence bias. First, and similarly to our previous study (Lebreton et al., 2018), we used both a free confidence elicitation method (experiment 1) and an incentivized method (experiment 2) and clearly replicate our results across these two methods. This indicates that the confidence bias cannot be attributed to the confidence elicitation mechanism. This is also supported by the fact that the confidence bias is observed despite the incentives in the primary task (gain and loss) being orthogonalized from the ones used to elicit confidence judgments (always framed as a gain). Second, an interesting feature of the present experiments is that, in contrast to our previous study (Lebreton et al., 2018), monetary outcomes are displayed after –rather than before-confidence judgments. At the time of decision and confidence judgments, the value of decision-contexts is implicitly inferred by participants and not explicitly displayed on the screen. Combined with the fact that loss and gain conditions were interleaved and that previous studies indicate that in a similar paradigm subjects remain largely unaware of the contextual manipulations (Bavard et al., 2018), this suggests that the biasing effect of the valence of monetary outcomes demonstrated in previous reports (Lebreton et al., 2018) is not due to a simple framing effect, created by the display of monetary gains or losses prior to confidence judgments.

We offer two interpretations for the observed effects of gains versus losses on confidence. In the first interpretation, we propose that loss prospects simply bias confidence downward. In the second interpretation, we propose that loss prospects improve confidence calibration over gain prospects, thereby correcting overconfidence. Following the first interpretation, the apparent improvement in confidence calibration observed in our study does not correspond to a confidence judgment improvement *per se*, but is a mere consequence of participants being overconfident in this task. Accordingly, in a hypothetical task where participants would be underconfident in the gain domain, while the loss prospects would aggravate this underconfidence under the first interpretation, they would improve confidence calibration (hence correct this underconfidence) under the second interpretation. Future research is needed to distinguish between the two potential mechanisms.

Regardless of the interpretation of the reported effects, we showed that confidence can be modelled as a simple linear and additive combination of three variables: previous confidence rating, choice difficulty and the context value inferred from the context-dependent reinforcement learning model. The critical contribution of the present study is the demonstration that confidence judgments are affected by the value of the decision-context, also referred to as context value. The context value is a subjective estimate of the average expected-value of a pair of stimuli: in our experimental paradigm, the context value is therefore neutral (equal to 0) at the beginning of learning, and gradually becomes positive in the reward-seeking conditions and negative in the punishment-avoidance conditions (Palminteri et al., 2015). The fact that the context-value significantly contributes to confidence judgments therefore complements our model-free results showing that outcome valence impacts confidence, while embedding it in the learning dynamics. The fact that the context value is a significant predictor of confidence judgments also suggests that context-dependency in reinforcement learning is not only critical to account for choice patterns but also to account for additional behavioral manifestations, such as confidence judgments and reaction times. This result therefore provides additional support for the idea that context values are explicitly represented during learning (Palminteri et al., 2015). Crucially, context-dependency has been shown to display locally adaptive (i.e. successful punishment-avoidance in the learning test) and globally maladaptive (i.e. irrational preferences in the transfer test) effects (Bavard et al., 2018). Whether the context-dependence of confidence judgments is adaptive or maladaptive remains to be elucidated and will require teasing apart the different interpretation of this effect discussed above.

Our findings are also consistent with a growing literature showing that in value-based decision-making, choice-difficulty, as proxyed by the absolute difference in expected subjective value between the available (Lebreton et al., 2009; Milosavljevic et al., 2010; Shenhav et al., 2014) is a significant predictor of confidence judgments (De Martino et al., 2013; Folke et al., 2016). Finally, the notion that confidence judgments expressed in preceding trials could inform confidence expressed in subsequent trails is relatively recent, but has received both theoretical and experimental support (Navajas et al., 2016; Rahnev et al., 2015) and intuitively echoes findings of serial dependence in perceptual decisions (Fischer and Whitney, 2014). In interleaved experimental designs like ours, successive trials pertain to different learning contexts. Therefore, the significant serial dependence of confidence judgments revealed by our analyses captures a temporal stability of confidence, which is context-independent. This result is highly consistent with the findings reported in Rahnev and colleagues (2015), which show that serial dependence in confidence can even be observed between different tasks.

Overall, our results outline the importance of investigating confidence biases in reinforcement-learning. As outlined in the introduction, most sophisticated RL algorithms assume representation of uncertainty and/or strategy reliability estimates, which allow them to flexibly adjust learning strategies or to dynamically select among different learning strategies. Yet, despite their fundamental importance in learning, these uncertainty estimates have, so far, mostly emerged as latent variables, computed from individuals’ choices under strong computational assumptions (Behrens et al., 2007; Collins and Koechlin, 2012; Daw et al., 2005; Donoso et al., 2014; Iglesias et al., 2013; Lee et al., 2014; Vinckier et al., 2016). In the present paper we propose that confidence judgments could be a useful experimental proxy for such estimates in RL. Confidence judgments indeed possess important properties, which suggest that they might be an important variable mitigating learning and decision-making strategies. First, confidence judgments accurately track the probability of being correct in stochastic environments, integrating expected and unexpected uncertainty in a close-to-optimal fashion (Heilbron and Meyniel, 2018; Meyniel et al., 2015a). Second, subjective confidence in one’s choices impacts subsequent decision processes (Braun et al., 2018) and information seeking strategies (Desender et al., 2018). Finally, confidence acts as a common currency and therefore can be used to trade-off between different strategies (de Gardelle and Mamassian, 2014; de Gardelle et al., 2016). With this in mind, biases of confidence could have critical consequences on reinforcement learning and reveal important features about the flexibility of learning and decision-making processes in different contexts. For instance, lower confidence in the loss domain – as demonstrated in the present report - could play an adaptive function, by allowing rapid behavioral adjustments under threat.

## Material and Methods

### Subjects

All studies were approved by the local Ethics Committee of the Center for Research in Experimental Economics and political Decision-making (CREED), at the University of Amsterdam. All subjects gave informed consent prior to partaking in the study. The subjects were recruited from the laboratory’s participant database (www.creedexperiment.nl). A total of 36 subjects took part in this study: 18 took part in experiment 1 (8/10 MF, age = 24.6±8.5), 18 in experiment 2 (8/10 MF, age = 24.6±4.3). They were compensated with a combination of a base amount (5€), and additional gains and/or losses depending on their performance during the learning task: experiment 1 had an exchange rate of 1 (in-game euros = payout); experiment 2 had an exchange rate of 0.3 (in game euros = 0.3 payout euros). In addition, in experiment 2, three trials (one per session) were randomly selected for a potential 5 euros bonus each, attributed based on the confidence incentivization scheme (see below).

### Power analysis and sample size determination

Power analysis were performed with GPower.3.1.9.2 (Faul et al., 2007). The sample size was determined prior to the start of the experiments based on the effects of incentives on confidence judgments in Lebreton et al. (2018). Cohen’s d was estimated from a GLM d = .941 t_23_ = 4.61, P = 1.23e-4). For a similar within-subject design, a sample of N=17 subjects was required to reach a power of 95% with a two-tailed one-sample t-test.

### Learning task

All tasks were implemented using MatlabR2015a^®^ (MathWorks) and the COGENT toolbox (http://www.vislab.ucl.ac.uk/cogent.php). In both experiments, the main learning task was adapted from a probabilistic instrumental learning task used in a previous study (Palminteri et al., 2015). Invited participants were first provided with written instructions, which were reformulated orally if necessary. They were explained that the aim of the task was to maximize their payoff and that gain seeking and loss avoidance were equally important. In each of the three learning session, participants repeatedly faced four pairs of cues - taken from Agathodaimon alphabet. The four cue pairs corresponded to four conditions, and were presented 24 times in a pseudo-randomized and unpredictable manner to the subject (intermixed design). Of the four conditions, two corresponded to reward conditions, and two to loss conditions. Within each pair, and depending on the condition, the two cues of a pair were associated with two possible outcomes (1€/0€ for the gain and −1€/0€ for the loss conditions in Exp. 1; 1€/0.1€ for the gain and −1€/−0.1€ for the loss conditions in Exp. 2) with reciprocal (but independent) probabilities (75%/25% and 25%/75%).

The reason for replacing the neutral outcome (0 euro) with a 10c gain or loss in Experiment 2 was to neutralize an experimental asymmetry between the gain and loss conditions, present in Experiment 1, which could have contributed to the valence impact on confidence in the partial information condition: when learning to avoid losses, subjects increasingly selected the symbol associated with a neutral outcome (0 euro), hence were provided more often with this ambiguous feedback^1^. It is worth noting that this asymmetry was almost absent in the complete feedback case where the context value can be inferred in both gains and losses thanks to the counterfactual feedback (e.g. a forgone loss), and nonetheless showed lower reported confidence. Yet, replacing the ambiguous neutral option with small monetary gains and losses in experiment 2 completely corrected the imbalance between the partial information gain and loss conditions.

At each trial, participants first viewed a central fixation cross (500-1500ms). Then, the two cues of a pair were presented on each side of this central cross. Note that the side in which a given cue of a pair was presented (left or right of a central fixation cross) was pseudo-randomized, such as a given cue was presented an equal number of times on the left and the right of the screen. Subjects were required to select between the two cues by pressing the left or right arrow on the computer keyboard, within a 3000ms time window. After the choice window, a red pointer appeared below the selected cue for 500ms. Subsequently, participants were asked to indicate how confident they were in their choice. In Experiment 1, confidence ratings were simply given on a rating scale without any additional incentivization. To perform this rating, they could move a cursor –which appeared at a random position-to the left or to the right using the left and right arrows, and validate their final answer with the spacebar. This rating step was self-paced. Finally, an outcome screen displayed the outcome associated with the selected cue, accompanied with the outcome of the unselected cue if the pair was associated with a complete-feedback condition.

### Matching probability and incentivization

In Experiment 2, participant’s reports of confidence were incentivized via a matching probability procedure that is based on the Becker-DeGroot-Marshak (BDM) auction (Becker et al., 1964) Specifically, participants were asked to report as their confidence judgment their estimated probability (p) of having selected the symbol with the higher average value, (i.e. the symbol offering a 75% chance of gain (G75) in the gain conditions, and the symbol offering a 25% chance of loss (L_25_) in the loss conditions) on a scale between 50% and 100%. A random mechanism, which draws a number (r) in the interval [0.5 1], is then implemented to select whether the subject will be paid an additional bonus of 5 euros as follows: If p ≥ r, the selection of the correct symbol will lead to a bonus payment; if p < r, a lottery will determine whether an additional bonus is won. This lottery offers a payout of 5 euros with probability r and 0 with probability 1-r. This procedure has been shown to incentivize participants to truthfully report their true confidence regardless of risk preferences (Hollard et al., 2015; Karni, 2009).

Participants were trained on this lottery mechanism and informed that up to 15 euros could be won and added to their final payment via the MP mechanism applied on one randomly chosen trial at the end of each learning session (3×5 euros). Therefore, the MP mechanism screens (Figure 3.A) were not displayed during the learning sessions.

### Transfer task

The 8 abstract stimuli (2×4 pairs) used in the third (i.e. last) session were re-used in the transfer task. All possible pair-wise combinations of these 8 stimuli (excluding pairs formed by two identical stimuli) were presented 4 times, leading to a total of 112 trials (Frank et al., 2004; Klein et al., 2017; Palminteri et al., 2015; Wimmer and Shohamy, 2012). For each newly formed pair, participants had to indicate the option which they believed had the highest value, by selecting either the left or right option via button press in a manner equivalent to the learning task. Although this task was not incentivized, which was clearly explained to participants, they were nonetheless encouraged to respond as if money was at stake. In order to prevent explicit memorizing strategies, participants were not informed that they would have performed this task until the end of the fourth (last) session of the learning test.

### Model-free statistics

All model-free statistical analyses were performed using Matlab R2015a. All reported p-values correspond to two-sided tests. T-tests refer to a one sample t-test when comparing experimental data to a reference value (e.g. chance: 0.5), and paired t-tests when comparing experimental data from different conditions. ANOVA are repeated measure ANOVAs.

### Computational modelling

#### Reinforcement-learning model

The approach for the reinforcement-learning modelling is identical to the one followed in Palminteri and colleagues (2015). Briefly, we adapted two models inspired from classical reinforcement learning algorithms (Sutton and Barto, 1998): the ABSOLUTE and the RELATIVE model. In the ABSOLUTE model, the values of available options are learned in a context-independent fashion. In the RELATIVE models, however, the values of available options are learned in a context-independent fashion.

In the ABSOLUTE model, at each trial t, the chosen (c) option value of the current context s is updated with the Rescorla-Wagner rule (Rescorla and Wagner, 1972):

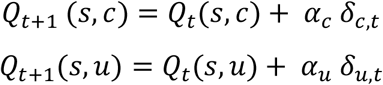

Where *α_c_* is the learning rate for the chosen (c) option and *α_u_* the learning rate for the unchosen (u) option, i.e. the counterfactual learning rate. *δ_c_* and *δ_u_* are prediction error terms calculated as follows:

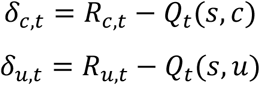

*δ_c_* is updated in both partial and complete feedback contexts and *δ_u_* is updated in the complete feedback context only.

In the RELATIVE model, a choice context value (*V*(*s*)) is also learned and used as the reference point to which an outcome should be compared before updating option values.

Context value is also learned via a delta rule:

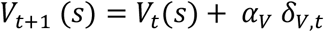

Where *α_v_* is the context value learning rate and *δ_v_* is a prediction error-term calculated as follows: if a counterfactual outcome *R_U,t_* is provided

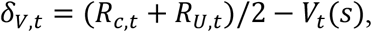

If a counterfactual outcome *R_U,t_* is not, provided, its value is replaced by its expected value *Q_t_*(*s,u*), hence

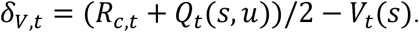

The learned context values are then used to center the prediction-errors, as follow:

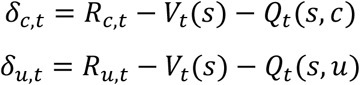

In both models, the choice rule was implemented as a softmax function: *P_t_*(*s,a*) = (1 + exp (*β*(*Q_t_*(*s,b*) − *Q_t_*(*s, a*))))^−1^, where *β* is the inverse temperature parameter.

#### Model fitting

Model parameters were estimated by finding the values which minimized the negative log likelihood (LLmax) and (in a separate optimization procedure) the negative log of posterior probability (LPP) of the observed choice given the considered model and parameter values. Note that the observed choices include both choices expressed during the learning test and choices observed during the transfer test, which were modelled using the option’s Q-values estimated at the end of learning. The parameter search was implemented using Matlab’s *fmincon* function, initialized at multiple starting points of the parameter space (Daw, 2011). Negative log-likelihoods (LLmax) were used to compute classical model selection criteria. The LPP was used to compute the exceedance probability and the expected frequencies of the model.

### Model comparison

We computed at the individual level (random effects) the Akaike’s information criterion (AIC),

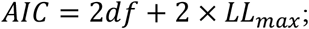

the Bayesian information criterion (BIC),

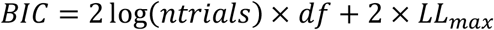

and the Laplace approximation to the model evidence (LPP);

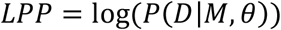

where *D, M* and *θ* represent the data, model and model parameters respectively. Following (Palminteri et al., 2015), *P*(*θ*) is calculated based on the parameters value retrieved from the parameter optimization procedure, assuming learning rates beta distributed (betapdf(parameter,1.1,1.1)) and softmax temperature gamma-distributed (gampdf(parameter,1.2,5)). Individual LPPs were fed to the mbb-vb-toolbox (https://code.google.com/p/mbb-vb-toolbox/) (Daunizeau et al., 2014). This procedure estimates the expected frequencies of the model (denoted PP) and the exceedance probability (denoted XP) for each model within a set of models, given the data gathered from all subjects. Expected frequency quantifies the posterior probability, i.e., the probability that the model generated the data for any randomly selected subject.

#### Confidence model

To model confidence ratings, we used the parameter and latent variables estimated from the best fitting Model (i.e. the RELATIVE model) under the LPP maximization procedure. Note that for Experiment 1, confidence ratings were linearly transformed from 1:10 to 50:100%.

Following the approach taken with the RL models, we designed two models of confidence: the FULL and the REDUCED confidence models.

In the FULL confidence model, confidence ratings at each trial *t*(*c_t_*) were modelled as a linear combination of the choice difficulty –proxied by the absolute difference between the two options expected value (*dQ_t_*), the learned context value (*V_t_*), and the confidence expressed at the preceding trial (*c*_*t*−1_).

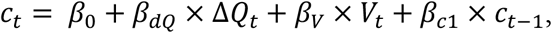

where

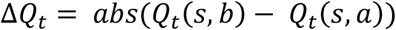

and *β*_0_, *β_dQ_*, *β_V_* and *β*_*c*1_ represents the linear coefficients of regression to be estimated.

In the REDUCED confidence model, we omitted the learned context value (*V_t_*), leading to

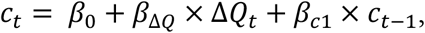

Those models were encapsulated in a generalized linear mixed-effect (glme) model, including subject level random effects (intercepts and slopes for all predictor variables). The model was estimated using Matlab’s *fitglme* function, which maximize the maximum likelihood of observed data under the model, using the Laplace approximation.

Modelled confidence ratings (i.e. confidence model fits) were estimated using Matlab’s *predict* function.

Because the REDUCED model is nested in the FULL model, a likelihood ratio test can be performed to assess whether the FULL model gives a better account of the data, while being penalized for its additional degrees-of-freedom (i.e. higher complexity). This test was performed using Matlab’s *compare* function.

To assess the specificity of V(s) we run two additional glmes including Σ*Q_t_* = *Q_t_*(*s,b*) + *Q_t_*(*s,a*) and the reaction time, respectively as model-based and model-free variables affected by the valence factor. We tested whether in these glmes V(s) still predicted confidence rating despite sharing common variance with these variables.

## Acknowledgment and funding

This work was supported by startup funds from the Amsterdam School of Economics, awarded to JBE. ML is supported by an NWO Veni Fellowship (Grant 451-15-015), and the Bettencourt Schueller Fondation. SP is supported by an ATIP-Avenir grant (R16069JS), the Programme Emergence(s) de la Ville de Paris, the Fyssen foundation and a Collaborative Research in Computational Neuroscience ANR-NSF grant (ANR-16-NEUC-0004). The Institut d’Etude de la Cognition is supported financially by the LabEx IEC (ANR-10-LABX-0087 IEC) and the IDEX PSL* (ANR-10-IDEX-0001-02 PSL*).

All data needed to evaluate the conclusions in the paper are present in the paper and/or the Supplementary Materials. Additional data related to this paper may be requested from the authors.

## Authors contributions

Designed the study: ML, SP and JBE. Collected the data: KB; Analyzed the data: ML. Interpreted the results: ML, SP and JBE. Drafted the manuscript: ML. Edited and finalized the manuscript: ML, SP and JBE.

## Footnotes

1 Note that, despite this asymmetry, there was no detectable difference in performance between gain and loss performance in the partial information in the Experiment 1.

